# Epigenetic inheritance and the evolution of infectious diseases

**DOI:** 10.1101/2020.08.29.273326

**Authors:** David V. McLeod, Geoff Wild, Francisco Úbeda

**Affiliations:** Centre D’Ecologie Fonctionnelle & Evolutive, CNRS, 34090 Montpellier, France; Institute of Integrative Biology, ETH Zürich, 8092 Zürich, Switzerland; Department of Applied Mathematics, The University of Western Ontario, London, Ontario N6A 5B7, Canada; Department of Biology, Royal Holloway University of London, Egham, Surrey TW20 0EX, United Kingdom

**Keywords:** epidemiology, evolutionary ecology, game theory, genomic imprinting, infectious diseases, mathematical model, measles

## Abstract

Genes with identical DNA sequences may show differential expression because of epigenetic marks. These marks in pathogens are key to their virulence and are being evaluated as targets for medical treatment. Where epigenetic marks were created in response to past conditions (epigenetically inherited), they represent a form of memory, the impact of which has not been considered in the evolution of infectious diseases. We fill this gap by exploring the evolution of virulence in pathogens that inherit epigenetic information on the sex of their previous host. We show that memories of past hosts can also provide clues about the sex of present and future hosts when women and men differ in their immunity to infection and/or their interactions with the sexes. These biological and social differences between the sexes are pervasive in humans. We show that natural selection can favour the evolution of greater virulence in infections originating from one sex. Furthermore, natural selection can favour the evolution of greater virulence in infections across sexes (or within sexes). Our results explain certain patterns of virulence in diseases like measles, chickenpox and polio that have puzzled medical researchers for decades. In particular, they address why girls infected by boys (or boys infected by girls) are more likely to die from the infection than girls infected by girls (or boys infected by boys). We propose epigenetic therapies to treat infections by tampering with the memories of infecting pathogens. Counterintuitively, we predict that successful therapies should target pathogen’s genes that inhibit virulence, rather than those enhancing virulence. Our findings imply that pathogens can carry memories of past environments other than sex (e.g. those related to socioeconomic status) that may condition their virulence and could signify an important new direction in personalised medicine.

## Introduction

In general, the term *epigenetics* refers to the differential expression of genes with identical DNA sequences in response to environmental factors ^17,19^—e.g. parental origin, stress level, social status. These environmental factors leave transient marks that do not change the DNA sequence of the gene, for example DNA methylation or histone modification ^7,17,19^. Some of these *epigenetic marks* are maintained during the lifetime of an individual (*epigenetically-acquired*) but they are not inherited from one generation to the next. Other marks however, are inherited across generations (*epigenetically-inherited*) ^17,19^. An example of epigenetic inheritance is the methylation of the promoter region of gene IGF2 in women’s but not in men’s germline ^16,18^. This epigenetic mark is transmitted across generations resulting in the differential expression of the maternally- and paternally-inherited copies of gene IGF2 in both sexes ^16,18^. Epigenetic inheritance thus opens up the fascinating possibility that genes have *memories* of environmental factors that they were exposed to in the past but are not exposed to in the present.

Due to its developmental and medical implications, epigenetic inheritance has received a lot of attention that, in turn, has advanced our knowledge on phenomena like genomic imprinting or cancer epigenetics ^13,30^. Despite the abundant attention that epigenetic inheritance has received, memories have rarely been considered when studying the epidemiology and evolution of infectious diseases. The oversight in exploring this possibility is not justified, as empirical evidence shows that epigenetic marks modify the expression of genes in pathogens controlling key functions like transmission or virulence ^7,8,11,22,26,32,34,37^. Here we work towards filling this gap.

We advance evolutionary theory by studying the evolution of virulence in pathogens that can remember the sex of the host they came from (henceforth *origin-specific virulence*). We outline conditions under which natural selection favours pathogens that retain the memory of their host of origin over those that do not. Furthermore, we make testable predictions about when pathogens will evolve to be more virulent when inherited from one sex as opposed to the other. In addition, we investigate when pathogens that can condition their virulence on the sex of their previous host (e.g., through epigenetic inheritance) will be selected to add information regarding the sex of their current host (e.g., through epigenetic plasticity) (henceforth *origin-&-sex-specific virulence*). We make explicit predictions about when pathogens will evolve to be more virulent when received from and infecting the same-sex as opposed to received from and infecting the opposite-sex.

Our results can explain puzzling observations of virulence patterns in measles, chickenpox and polio wherein those with infections contracted from same-sex individuals develop a less virulent infection compared to those infected by the opposite sex ^2,25,28,31^. That in developing countries girls infected with measles by boys are more likely to die from the infection that girls infected by girls, is a well established result but remains poorly understood after more than two decades of research ^2,28^. What is puzzling about this pattern is that it cannot be explained by differences in virulence between girls and boys (i.e. due to differences between the sexes in their immune system) as differences between the sexes will affect equally infections received from girls and boys. Here we argue that pathogens with epigenetic memories of the host from which they came, can explain the complex patterns of virulence found in measles, chickenpox and polio. Finally, we explore the implications of our model for the treatment of infectious diseases. In particular, we predict when the use of drugs that erase epigenetically inherited marks in pathogens will reduce their virulence.

## Results

### Model formulation and analysis

We advance existing models ^5,15^ by allowing epigenetic information on the type of host the pathogen originated from (see Methods for details).

We place all individuals who either host the pathogen – or could possibly host the pathogen – in one of two categories. Category *j* = *f* individuals are female, and category *j* = *m* individuals are male. Individuals of a given sex-*j* (henceforth *sex*) infected with a pathogen that most recently originated from a sex-*k* host (henceforth *origin*), recovers (becoming susceptible again), dies from causes unrelated to the infection at rate *µ*_*j*_ (*natural mortality*), or dies from causes related to the infection at a rate *α*_*j,k*_ (*virulence*).

We make the standard assumption that there is a trade-off between virulence and transmission ^4^. In particular, the greater the replication rate of a pathogen the higher the mortality rate of hosts due to the disease and the higher the transmission rate of the pathogen to new hosts. If *β*_*i←*(*j,k*)_ denotes the transmissibility of a (*j, k*) infection to a susceptible sex-*i* host, then in mathematical terms *β*_*i←*(*j,k*)_ = *β*_*i←*(*j,k*)_(*α*_*j,k*_) and *dβ*_*i←*(*j,k*)_/*dα*_*j,k*_ *>* 0.

We use “*•*” to indicate restrictions associated with transmissibility. For example, *β*_*•←*(*j,k*)_ indicates dependence upon *j* and *k* but not *i*. To be clear, the use of “*•*” here only indicates a lack of direct dependence. For example, *β*_*•←*(*j,•*)_ may still depend on the sex of the host of origin indirectly through direct dependence on *α*_*j,k*_. Despite the possibly complicated relationship between transmissibility and virulence, when describing our results, we will want to compare various transmissibilities that arise using some benchmark level of virulence, denoted *α*_bm_. For simplicity, then, we will discuss only benchmarked *β*_*i←*(*j,k*)_s below. Consequently, *β*_*i←*(*j,k*)_ should be henceforth understood as *β*_*i←*(*j,k*)_(*α*_bm_).

In general, we allow different categories of host to belong to one of two subpopulations, *𝓁* = *A* and *𝓁* = *B*, and we assume incomplete mixing between them. Subpopulations differ with respect to the pattern of contact between host types and/or the influx susceptible hosts. We use 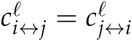 to denote the probability with which an *i, j* interaction occurs. Influx of new susceptible hosts may be biased in favour of one particular category of host, or one particular subpopulation and we use 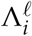 to denote the birth of sex-*i* hosts. If 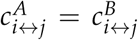 and 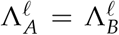 for all *i, j*, then the two subpopulation model collapses to a one-population model. We treat these two scenarios separately.

The focus of our research is on the evolution of epigenetically inherited virulence. In the first instance, we allow pathogens to adjust their virulence using epigenetically inherited information only. In our model, this translates into studying virulence that is constrained to evolve conditional upon the sex of the host from which the infection originated (origin-specific virulence, denoted *α*_*•,k*_). Later, we allow pathogens to adjust their virulence using both inherited and acquired information. In our model, this translates into studying virulence that is unconstrained (origin-&-sex-specific virulence, denoted *α*_*j,k*_). Our model also allows research into virulence in the absence of information and virulence adjusted by acquired information only. However, these last two forms of virulence have been studied in detail elsewhere ^10,36^.

In the Methods section we develop mathematical descriptions of the direction of evolution favoured by natural selection. In our model, the action of selection ultimately produces an *evolutionarily stable* (ES) ^9^ pattern of virulence, 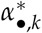 or 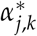 Using these mathematical descriptions we address two questions: i. When does origin-specific or origin-&-sex-specific virulence evolve? ii. When do we expect to observe greater virulence of pathogens with a particular origin or a particular origin in a particular sex? We pay attention to contrasting patterns of virulence exhibited by infections transmitted between same-sex individuals against patterns exhibited by infections transmitted between opposite-sex individuals. This contrast is deserving of special attention as it is related to complex patterns of virulence in measles, chickenpox and polio—patterns that can not be explained by recourse to sex-specific virulence.

### Origin-specific virulence

#### No population structure

Firstly, origin-specific virulence evolves when transmissibility is origin-specific, meaning *β*_*i←*(*j,k*)_ = *β*_*•←*(*•,k*)_.

In general, when there are origin-specific effects on transmission, the ES virulence of a pathogen inherited from one sex (denoted *k*) will be greater than that of the same pathogen inherited from the opposite sex (denoted *¬k*), when the transmissibility of the former is lower than the transmissibility of the latter (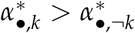 when *β*_*•←*(*•,k*)_ *< β*_*•←*(*•,¬k*)_; see Methods).

The intuitive reason is that a pathogen’s origin provides direct information on transmissibility. Virulence then evolves to be lower in the sex that transmits more readily as a result of the virulence-transmission trade-off.

Secondly, origin-specific virulence can sometimes evolve when transmissibility is sex-specific, but not always. Transmissibility can be considered to be sex-specific when it depends on the sex of the host in which the pathogen currently resides (*β*_*i←*(*j,k*)_ = *β*_*•←*(*j,•*)_), when it depends on the sex of the host in which the new infection will become established (*β*_*i←*(*j,k*)_ = *β*_*i←*(*•,•*)_), or when it depends on both the sex in which the pathogen currently resides and the sex that follows (*β*_*i←*(*j,k*)_ = *β*_*i←*(*j,•*)_). Of these three possibilities, only the third leads to origin specific virulence (see Methods).

Where sex-specific transmissibility supports the evolution of origin-specific virulence, it does so in two distinct ways. If the average same-sex transmissibility exceeds the average cross-sex transmissibility, then greater virulence occurs in infections originating from the sex with greater average infectivity (if 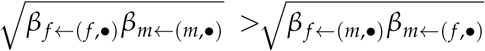, then 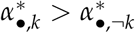 when 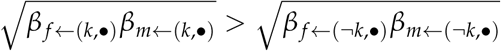; see Figure 1.a.i). By contrast, if average cross-sex transmissibility exceeds same-sex transmissibility, then the pattern is reversed: infections originating from the sex with the lower infectivity are more virulent (see Figure 1.a.ii).

**Figure 1a:**
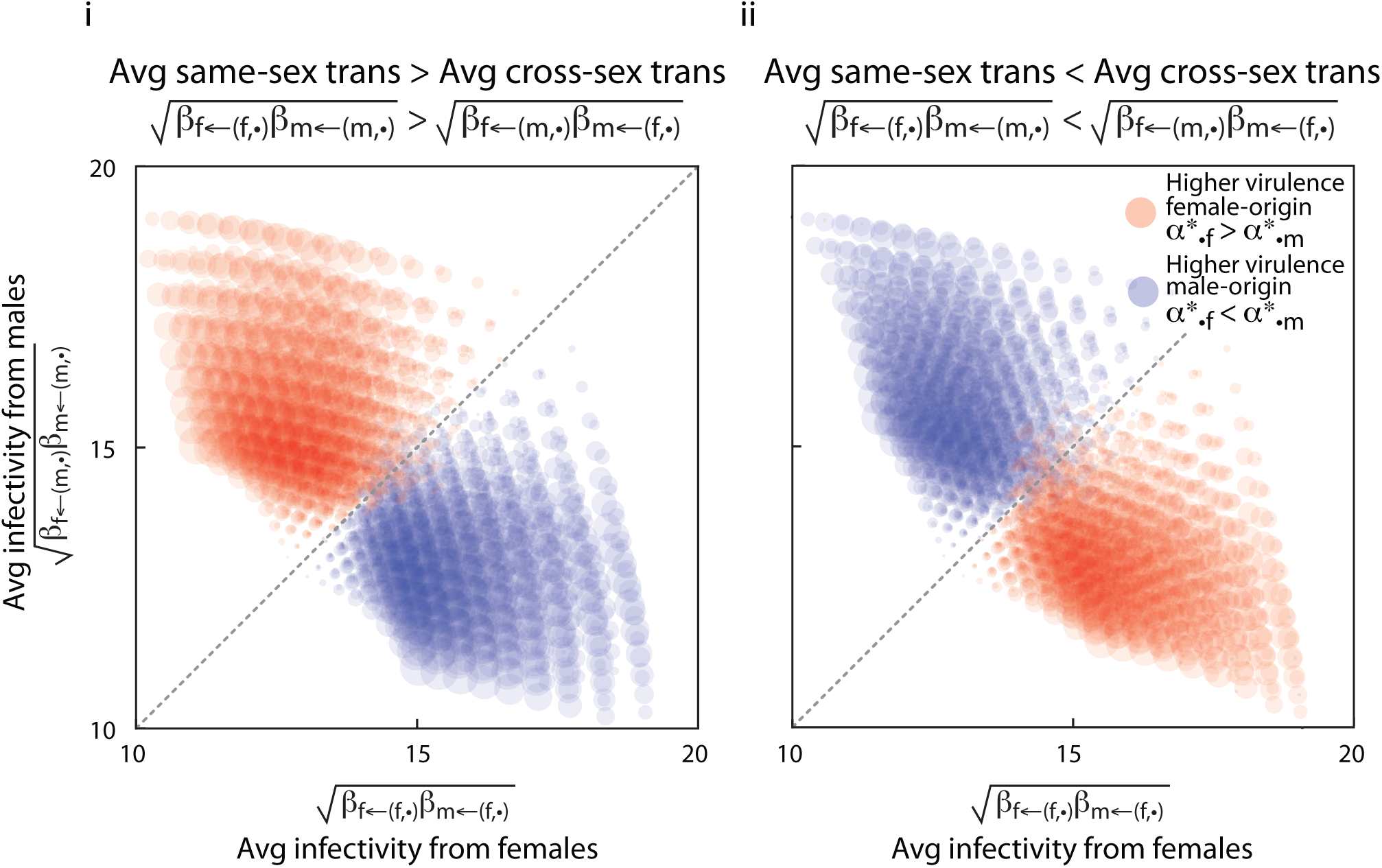
Evolution of origin-specific virulence with different infectivity and susceptibility between the sexes. Here pathogens inherit epigenetic marks that allow them to condition their virulence on the sex of the host they originated from. **Panel a: Numerical results** Shows the difference between evolutionarily stable virulence of female-acquired infections and male-acquired infections plotted as circles whose area scales with the extent of the difference itself. Circles are centred according to the average infectivity of females, defined as (*β*_*m←* (*f*, •) *β* *f←* (*f*, •)_)^1/2^, and average infectivity of males, defined as (*β* _*f←* (*m*, •)_ *β*_*m←* (*m*, •))_^1/2^. These results are based on a one-population model in which there are no sex-specific differences in background mortality, no possibility of recovery, and no difference in the influx of sexes to the population. Parameters (see Methods): *c-values all 1, m* = 35, *ρ* = 0.5, *µ* _*•*_ = 0.5, *γ* _*•*_ = 0, *θ*_*ij•*_ *ranged from 0*.*5 to 1*.*5 and all combinations were considered (*9^4^ *different combinations)*. **Panel i: Higher same-sex transmissibility**. Shows the results when the average transmissibility between individuals of the same sex, defined as (*β* _*f ←*(*f, •*)_ *β*_*m←*(*m,•*)_)^1/2^, is higher than the average transmissibility between individuals of the opposite sex, defined as (*β* _*f←* (*m*, •)_ *β*_*m←* (*f*, •)_)^1/2^ **Panel i i: Lower same-sex transmissibility**. Shows the results when the average transmissibility between individuals of the same sex is lower than the average transmissibility between individuals of the opposite sex.

**Figure 1:**
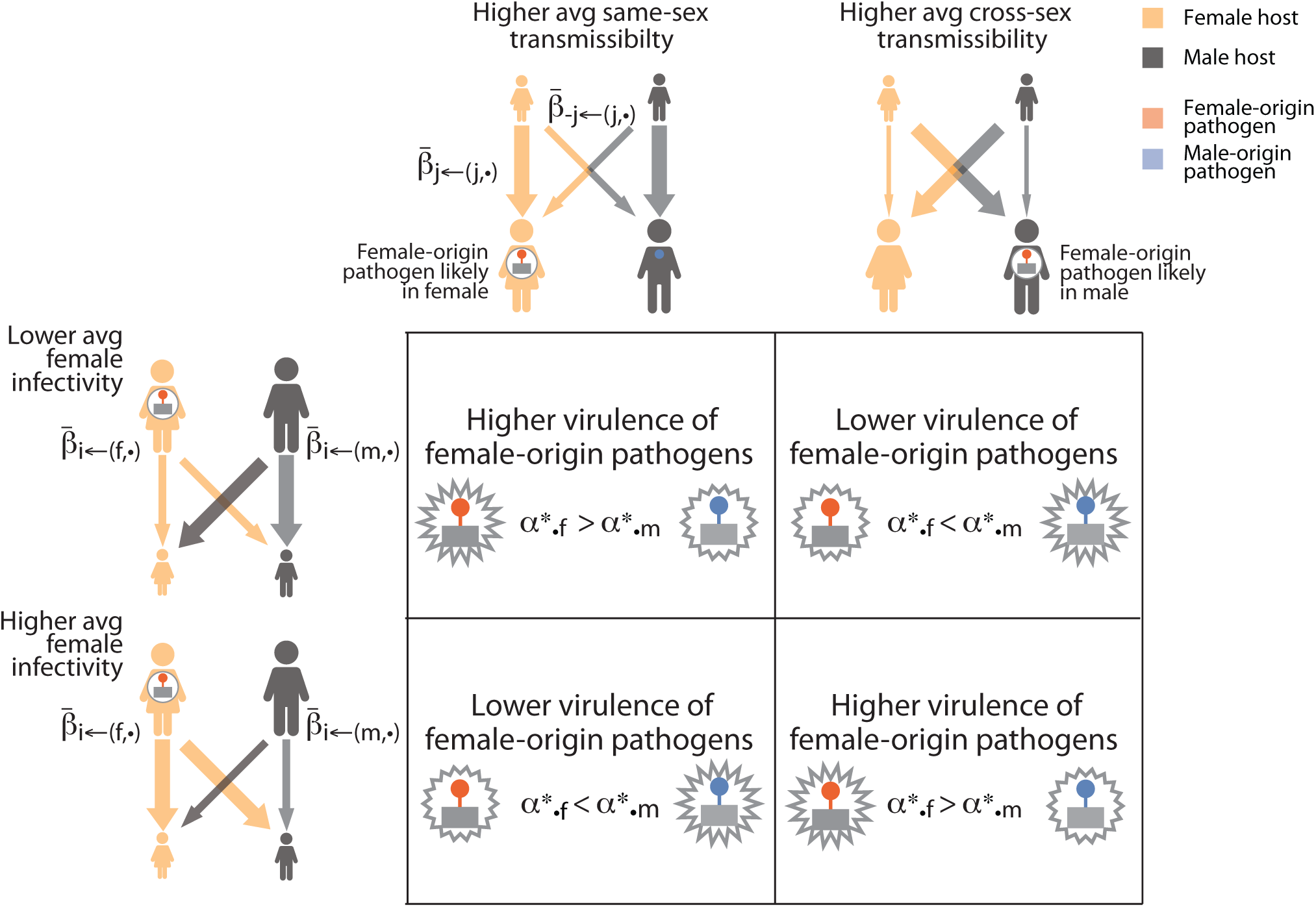
Evolution of origin-specific virulence with different infectivity and susceptibility between the sexes. Here pathogens inherit epigenetic marks that allow them to condition their virulence on the sex of the host they originated from. **Panel b: Intuitive explanation**. Provides intuitive explanations to the results in panels a.i and a.ii. Explanations are arranged as a table considering the four possible combinations of: higher or lower average same-sex transmissibility and higher or lower average female infectivity. For example, when average same-sex transmissibility is higher than average cross-sex transmissibility, a female-origin pathogen is more likely to be currently infecting a female host. When on average female hosts are more likely to transmit infections, pathogens evolve to be less virulent when they are of female-origin because they have a greater opportunity for transmission. Epigenetically inherited marks inform of the sex of the current host.

The intuition behind this result is that a pathogen’s origin provides information about the sex of the current host. Virulence then evolves to be lower in the sex that can infect more readily (Figure 1.b). For example, if the average cross-sex transmissibility is higher than the average same-sex transmissiblity, then a pathogen with female-origin is more likely to be infecting a male. In this case, when the average transmissibility from a sex (the *infectivity* of that sex) is higher in males, pathogens are selected to evolve lower virulence in males which corresponds to lower virulence when of female-origin (Figure 1.b).

#### Population structure

In this section we focus on one instance in which origin-specific virulence does not evolve in the absence of population structure. Suppose transmission depends only on the sex to which the pathogen is transmitted (*susceptibility* of a sex), *β*_*i←*(*j,k*)_ = *β*_*i←*(*•,•*)_. Here, origin-specific virulence evolves as long as subpopulations differ with respect to the sex-specific influx of new susceptible individuals, or with respect to contact structure (Figure 2.a and Methods). In particular, when sex-*k* is less susceptible to infection, the virulence of pathogens with sex-*k* origin evolves to be greater (if *β*_*k←*(*•,•*)_ *< β*_*¬k←*(*•,•*)_, then 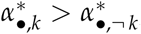; Figure 2.a).

**Figure 2:**
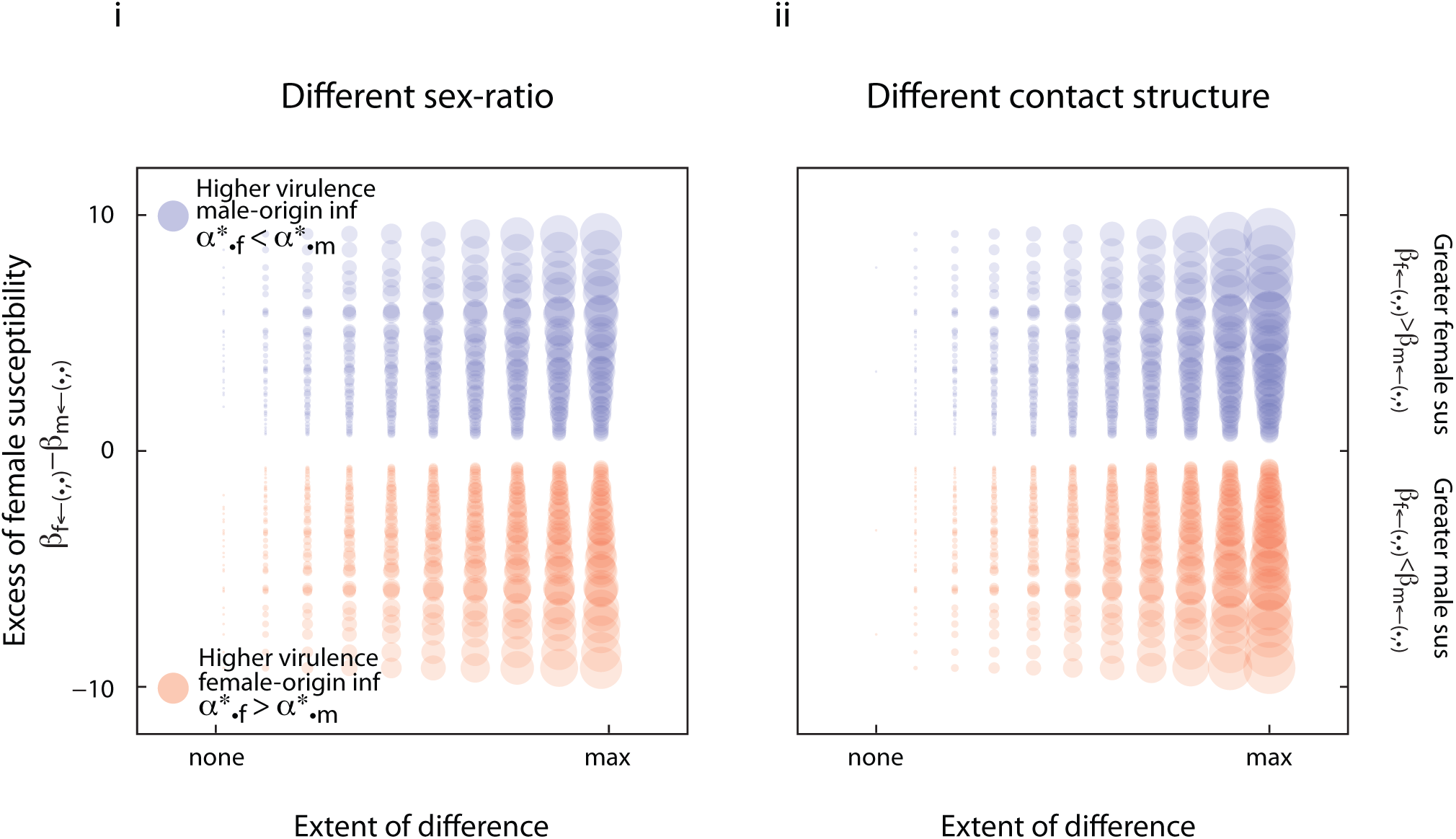
Evolution of origin-specific virulence with different sub-populations and different susceptibility between the sexes. Here pathogens inherit epigenetic marks that allow them to condition their virulence on the sex of the host they originated from. We considered that there are two-subpopulations (sub-populations A and B). **Panel a: Numerical results**. Shows the difference between evolutionarily stable virulence associated with female-acquired infections and male-acquired infections plotted as circles whose area scales with the extent of the difference itself. Circles are centred according to the extent of the difference between sub-populations and the excess female susceptibility, defined as *β* _*f←* (•,•)_ *β*_*m*← (•,•)_. These results assume that there are no sex-specific differences in background mortality, there was no possibility of recovery, and there was only weak mixing of populations (*σ* = 0.25). **Panel i: Sub-populations differ in their sex-ratio**. The difference between sub-populations in their influx of sexes varies continuously between a no-difference scenario to a maximal difference scenario in which influx to population B is strongly female biased while that to population A is strongly male biased (*ρ*_*A*_ *< ρ*_*B*_). We assume that sub-populations are identical with respect to their contact patterns 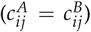. **Panel ii: Sub-populations differ in their contact structure**. The differences between sub-populations in their pattern of contact between sexes varies continuously from a no-difference scenario to a maximal-difference scenario in which population *A* contacts are exclusively same-sex in nature but population *B* contacts are exclusively cross-sex in nature 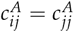 and 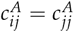).We assume that sub-populations experience equal and unbiased influx of the different sexes (*ρ*_*A*_ = *A ρ*_*B*=_= 0.5).

**Figure 2:**
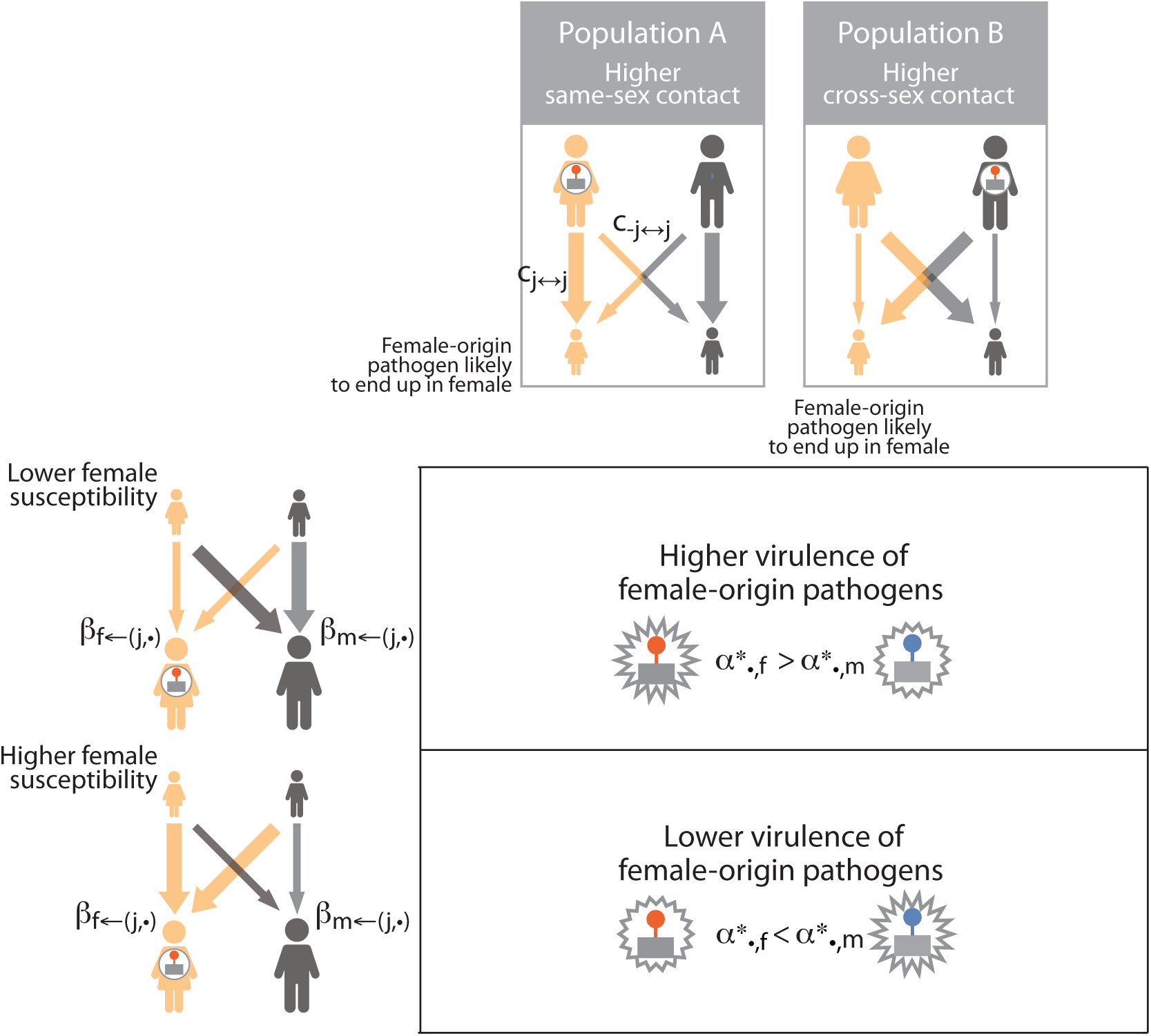
Evolution of origin-specific virulence with different sub-populations and different susceptibility between the sexes. Here pathogens inherit epigenetic marks that allow them to condition their virulence on the sex of the host they originated from. We considered that there are two-subpopulations (sub-populations A and B). **Panel b: Intuitive explanation**. Provides intuitive explanations to the results in panels a.i and a.ii. Explanations are arranged as a table considering the four possible combinations of: higher or lower average same-sex transmissibility and higher or lower average female infectivity. For example, when average same-sex transmissibility is higher than average cross-sex transmissibility, a female-origin pathogen is more likely to be currently infecting a female host. When on average female hosts are more likely to transmit infections, pathogens evolve to be less virulent when they are of female-origin because they have a greater opportunity for transmission. Epigenetically inherited marks inform of the sex of the current host.

The intuition behind this result is that the differences between sub-populations establishes a positive correlation between the sex of the host the pathogen will infect and the sex of the host from which the pathogen originated. Virulence evolves to be lower in the sex that can be infected more readily (Figure 2.b). When susceptibility differs between the sexes, pathogens evolve to be less virulent when they originated from the sex they are more likely to end up infecting, independently of the sex of the current host. For example, if subpopulation A shows a higher rate of same-sex contacts and subpopulation B a higher rate of cross-sex contacts, a female-origin pathogen in either of these populations is likely to end up infecting a female next. When the susceptibility is higher in males, pathogens are selected to evolve higher virulence when of female-origin (Figure 2.b).

### Origin-&-sex-specific virulence

So far we have explored the evolution of virulence when pathogens can only make use of inherited information. Now, we extend our analysis to include pathogens that can adjust their virulence in response to inherited information about the sex of the host they came from, and acquired information about the sex of the host in which they currently reside (we have defined this as origin-&-sex-specific virulence). This extension is motivated by the complex virulence patterns observed in measles, chickenpox and polio ^1,25^. These infections result in greater mortality in girls infected by boys and in boys infected by girls ^1,25^, a pattern that cannot be explained by invoking origin-specific or sex-specific virulence alone. Of course, sex-specific virulence could explain greater mortality arising from infections occurring in a given sex, and origin-specific virulence could explain greater mortality in infections originating from a given sex, but it takes origin-&-sex-specific virulence to explain the reported interaction effects.

#### No population structure

Origin-&-sex-specific virulence evolves when transmissibility depends on the sex of the host the pathogen originated from and the sex of the host the pathogen is currently found, that is *β*_*i←*(*j,k*)_ = *β*_*•←*(*j,k*)_. In particular, if transmissibility of a pathogen is greater when the sex of its current host matches the sex of the host from which the infection was acquired, then greater cross-sex virulence evolves (if *β*_*•←*(*k,k*)_ *> β*_*•←*(*k,¬ k*)_ then 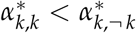).

A situation like this may arise when pathogens take time to adjust to current environments. If such an adjustment is required, the transfer from one sex to the other could delay disease progression and ultimately impact transmissibility in an origin-&-sex-specific way.

#### Population structure

The two subpopulations model, once again, expands the scope for the evolution of conditional expression of virulence. In this section, we explore two ways in which the expansion of scope can occur.

Firstly, we find that, in the presence of population structure, origin-&-sex-specific virulence can evolve even when transmissibility is the same for all host types (that is, *β*_*i←*(*j,k*)_ = *β*_*•←*(*•,•*)_). In this case, however, subpopulations must differ with respect to the sex-specific influx of susceptible hosts, and the pattern of contact between host types. Origin-&-sex-specific virulence is favoured, then, because it is an indirect response to the local conditions the pathogen is likely to encounter.

As an example, consider a case in which contacts in one sub-population tend to occur more frequently between same-sex individuals, whereas contacts in the other subpopulation tend to be between individuals of the opposite sex. Suppose further that opportunities to create new infections are more abundant in the former subpopulation because birth rates there are greater. Under these conditions, we find higher virulence from infections acquired from same-sex individuals on average, meaning 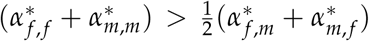, as shown in Figure 3. That figure also shows how the predictions change when assumptions about the birth rate and the pattern of contacts vary.

**Figure 3:**
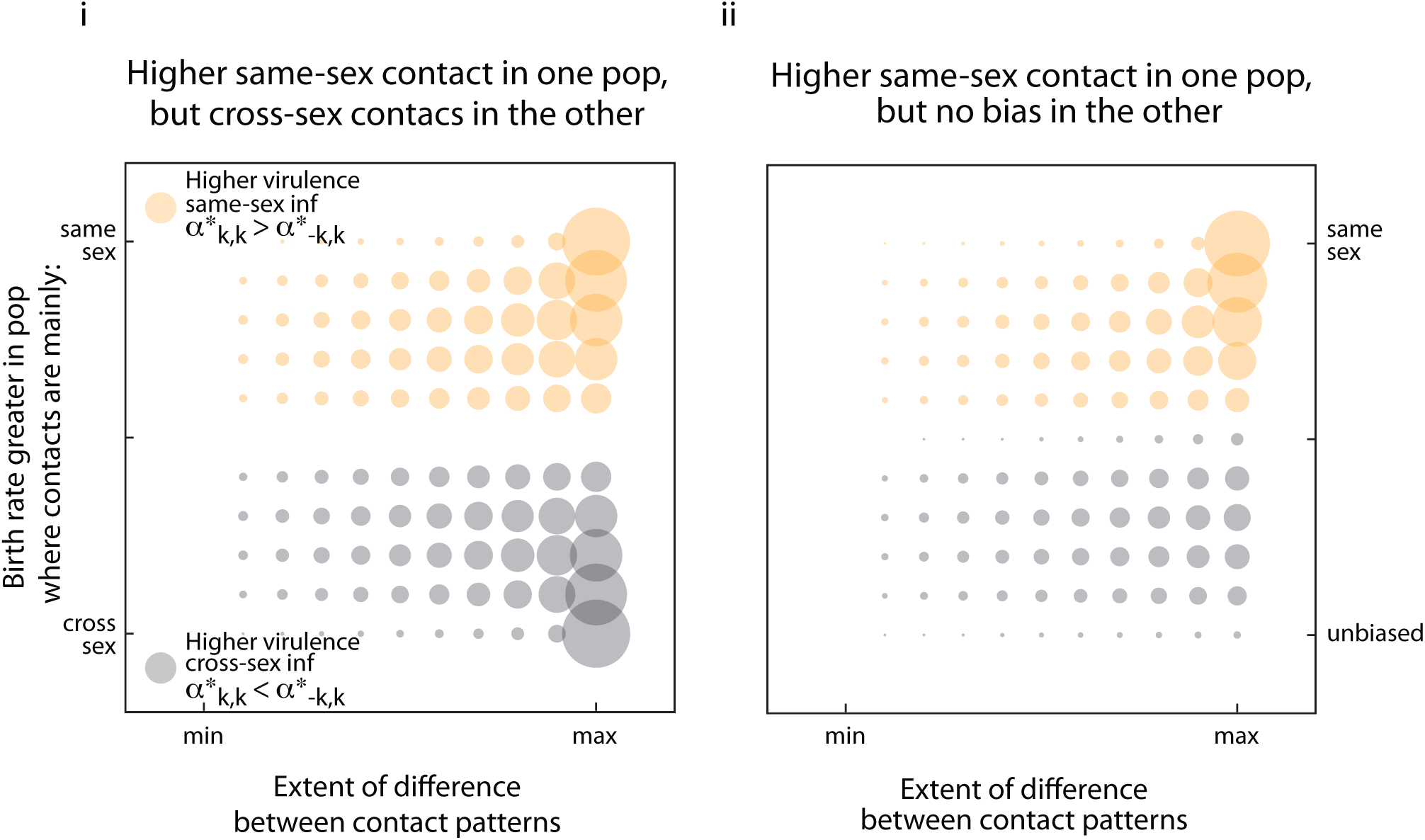
Evolution of origin-&-sex-specific virulence with sub-populations of different contact structure and different size. Here pathogens inherit epigenetic marks and use environmental cues that allow them to condition their virulence on the sex of the host they originated from and the sex of the current host. We assumed that there are two subpopulations. This figure shows the difference between average virulence associated with infections acquired from same-sex transmission, defined as 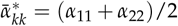, and those acquired from cross-sex transmission, defined 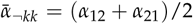, plotted here as circles whose area scales with the extent of the difference itself. Circles are centred according to the relationship between the contact patterns that exist in the respective sub-populations, and the extent of the bias in birth rate between subpopulations. These numerical results assume no sex-specific differences in transmissibility or susceptibility (*β*_•←•,•_), no sex-specific background mortality, no possibility of recovery, and weak mixing of populations (*σ* = 0.25). **Panel i: Higher same-sex contact in one population but lower in the other**. As the difference with respect to contact patterns between the populations increases, contacts in one sub-population become more strongly biased toward same-sex interactions while those in the other sub-population become equally biased but toward cross-sex interactions. Opportunities for the creation of new infections can be strongly associated with same-sex interactions when the influx of new susceptible hosts through birth strongly favours the population with that contact pattern. Birth rates may also be greater in the subpopulation associated with cross-sex interactions, in which case the negative consequences of pathogen virulence are lower there and greater cross-sex virulence evolves. **Panel ii: Higher same-sex contact in one population but no bias in the other**. Assumes no bias in the same-sex vs cross-sex pattern of contact in one of the two sub-populations. In this case the opportunity to create new infections ranges between same-sex bias and no bias as the influx of new susceptible hosts shifts through births from favouring one sub-population to favouring the other.

The intuition behind this result rests on a two-part rationale. First, when births balance deaths on a global scale, as they do in our model at equilibrium, a higher birth rate in one subpopulation disproportionately buffers the negative consequences of virulence there. As a result, there is less disincentive to be virulent in the population where same-sex contacts dominate. Second, knowing that the sex of the host of origin matched (resp. did not match) the sex of the current host would suggest to a pathogen that the consequences of virulence are less (resp. more) dire than they might be otherwise.

Secondly, the evolution of origin-&-sex-specific virulence can be supported in the presence of sex-specific transmissibilities as well. In order for this virulence pattern to evolve transmissibilities must show both sex-specific susceptibility and sex-specific infectivitity (*β*_*i←*(*j,k*)_ = *β*_*i←*(*j,•*)_). If transmissibilities are sex-specific, though, we can relax conditions on subpopulations, insisting that they only differ with respect *either* the sex-specific in susceptible hosts, *or* the pattern of contacts between host types.

In general when average same-sex transmissibility is higher, then infections on one sex originating from the opposite sex show greater virulence on average (if 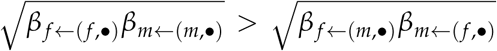, then 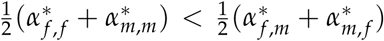; see Figure 4).

**Figure 4:**
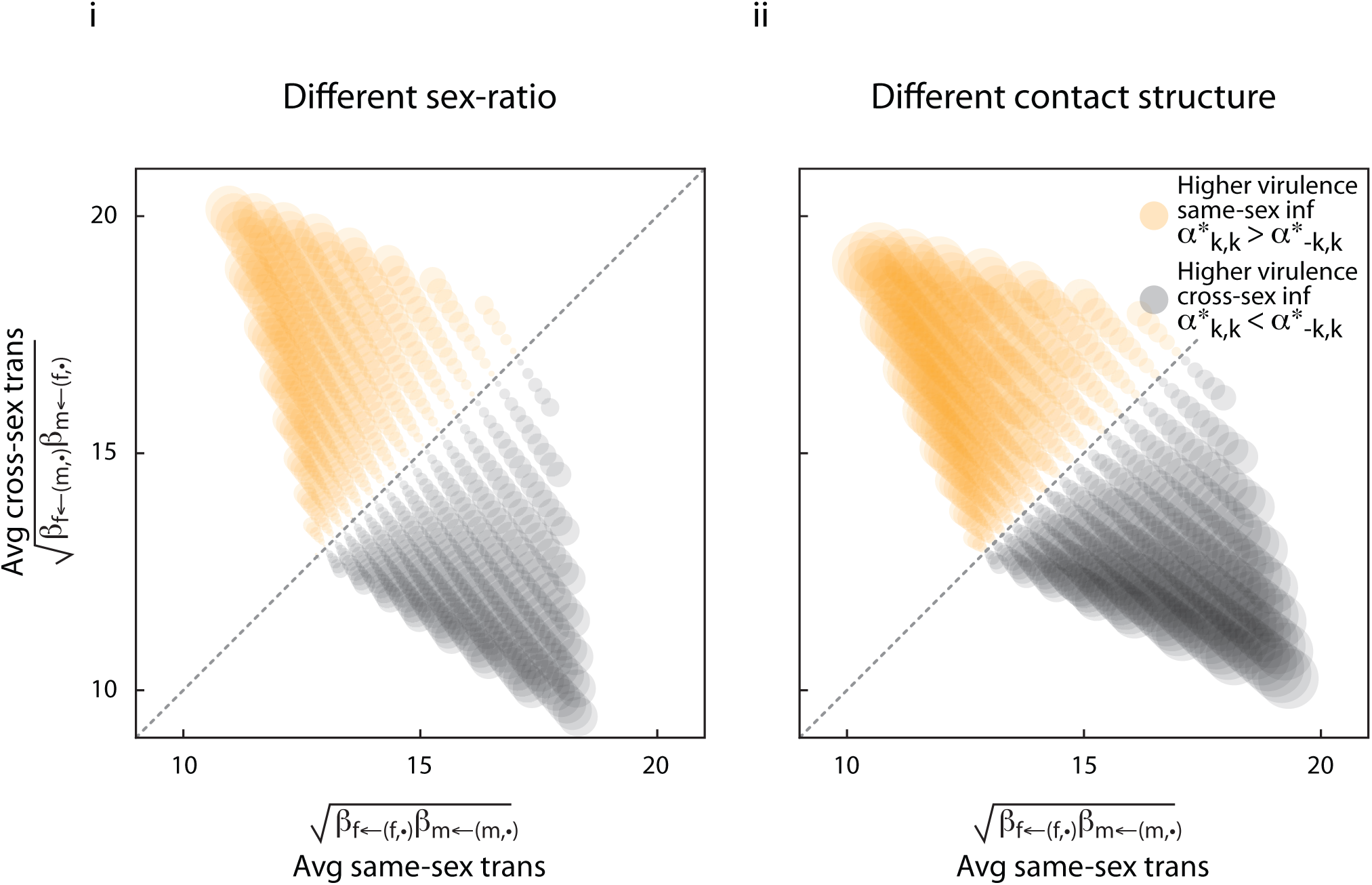
Evolution of origin-&-sex-specific virulence with different same-sex and cross-sex transmission and different sub-populations. Here pathogens inherit epigenetic marks and use environmental cues that allow them to condition their virulence on the sex of the host they originated from and the sex of the current host. We considered that there are two-subpopulations. This figure shows the difference between average virulence associated with infections acquired from same-sex transmission and those acquired from cross-sex transmission plotted here as circles whose area scales with the extent of the difference itself. Circles are centred according to average same-sex transmissibility and average cross-sex transmissibility. We assume no sex-specific differences in background mortality, no possibility of recovery, and weak mixing of populations (*σ* = 0.1). **Panel i: Sub-populations differ in their sex-ratio**. Influx of sexes to each sub-population is biased in exactly opposite directions (*ρ*_*A*_ = 1 *- ρ*_*B*_), but pattern of deliberate contact does not differ between populations 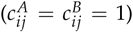. **Panel ii: Sub-populations differ in their contact structure**. Influx of sexes to each population is unbiased (*ρ*_*A*_ = *ρ*_*B*_ = 0.5), but pattern of deliberate contact differs between populations (same-sex contact in *A* is more common: 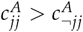, and cross-sex contact is more common in 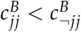.

## Discussion

Our research shows that genes in pathogens can be selected to retain epigenetic memories of the sex of the host from which they originated. With this information they may evolve greater virulence when they are inherited from a female rather than a male, or vice-versa. We have called this origin-specific virulence.

Pathogens are selected to retain epigenetic marks that allow for origin-specific virulence when these marks convey reliable information about: the sex of the host the pathogen was infecting (past), the sex of the host the pathogen is infecting (present), or the sex of the host the pathogen will be infecting (future) (Figure 5.a). To support the evolution of origin-specific virulence, the transmissibility in women and men and/or the interactions between the sexes must differ (see Figure 5.a and Methods for examples). These biological and social differences are pervasive in human populations: differences between the sexes in infectivity and in susceptibility are ubiquitous ^3,6,14,21,23,24,33^; differences between social groups in their sex-ratios and/or their contact pattern between individuals of the same and the opposite sexes have been widely reported ^27,35^.

**Figure 5:**
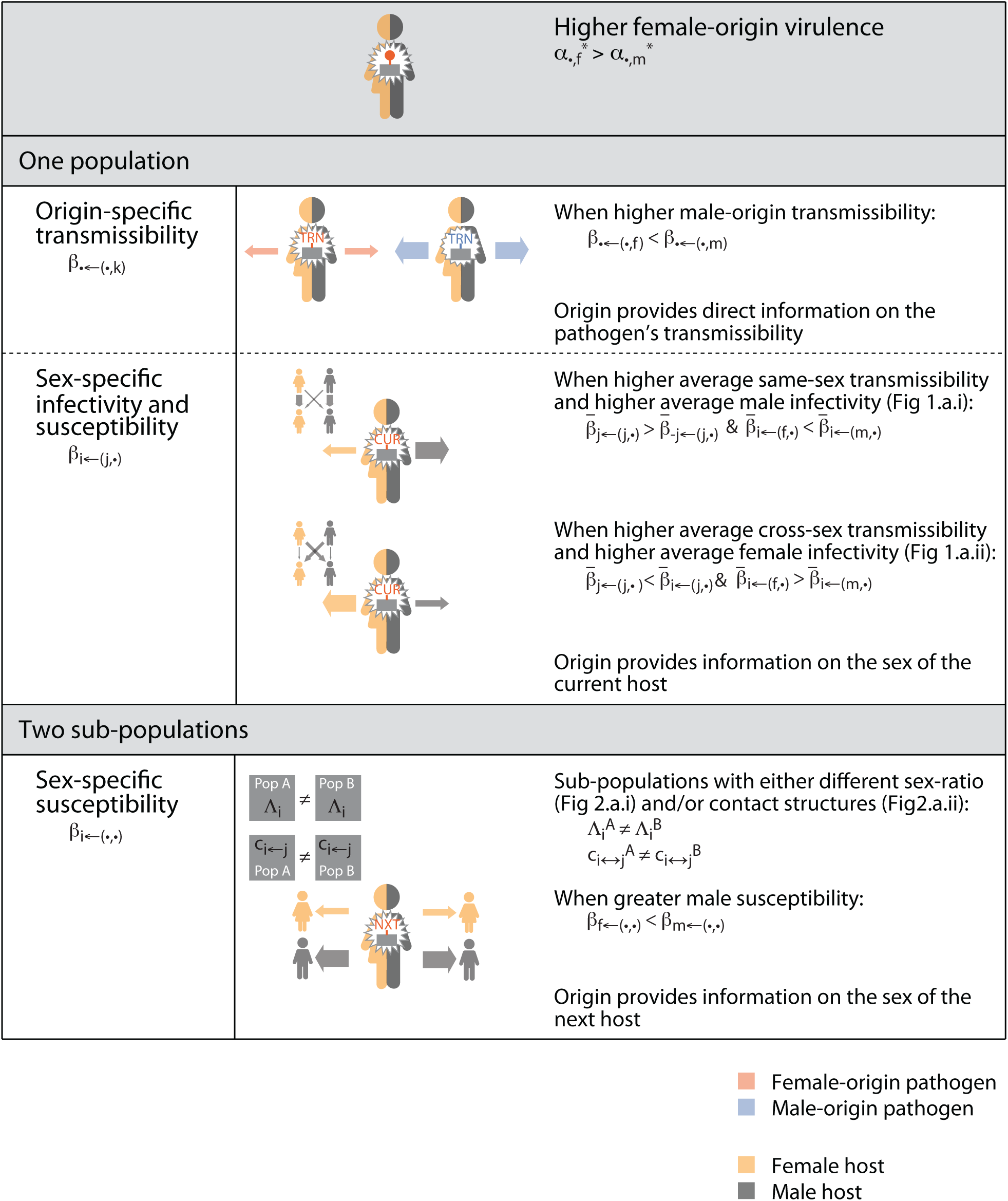
Summary of results. Panel i: Summary of results regarding the evolution of origin-specific virulence. We focus on the case when female-origin virulence will be greater than male-origin one. Each row indicates a set of different assumptions regarding the population structure and pathogen’s transmissibility.

**Figure 5:**
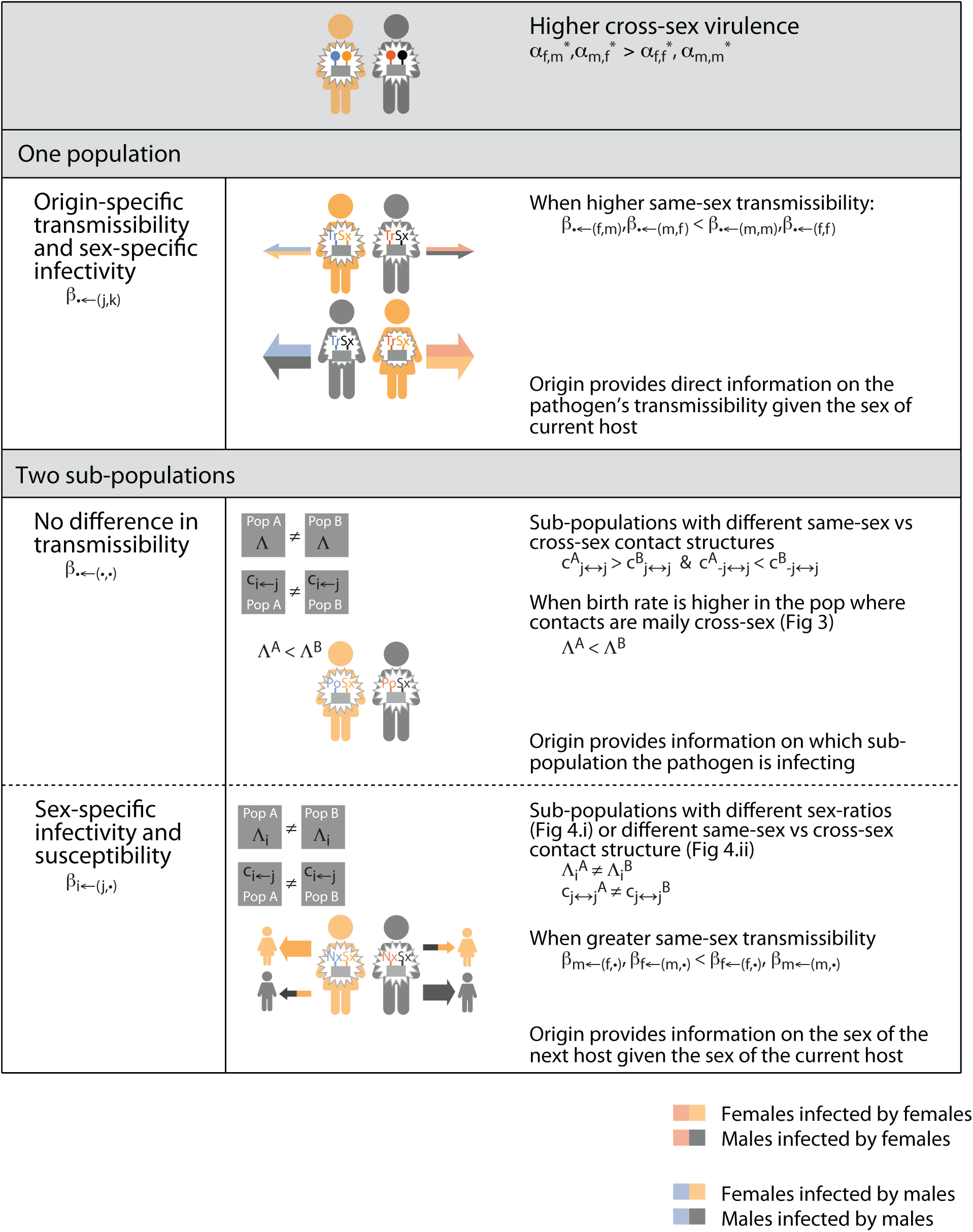
Summary of results. Panel ii: Summary of results regarding the evolution of origin-&-sex-specific virulence. We focus on the case when cross-sex virulence will be greater than same-sex one. Each row indicates a set of different assumptions regarding the population structure and pathogen’s transmissibility.

Our research shows that pathogens can be selected to acquire information (epigenetic or not) on the sex of the current host to existing epigenetically inherited information on the sex of the host from which they originated. In combination, both pieces of information open the door to the evolution of relatively complex patterns of virulence. We use the term origin-&-sex-specific virulence to refer to these complex patterns that, incidentally, cannot be explained by origin-specific virulence or sex-specific virulence alone.

Pathogens evolve origin-&-sex-specific virulence when inherited and acquired marks convey reliable information about: the sex of the host the pathogen was infecting (past), the sex of the host pathogen is infecting (present), or the sex of the host pathogen will be infecting (future) (Figure 5.b). Furthermore, when a population is structured into subpopulations that differ in a sex-specific way with respect to demographics and patterns of socialisation, then origin-&-sex marks can provide pathogens with crucial clues about their success (Figure 5.b and Methods for some examples).

Our findings provide a possible solution to the long-standing medical research on complex pattern of virulence in measles, chickenpox and polio. More than two decades ago, it was observed that, in countries with limited access to medical treatment, girls infected with measles by boys were 2.5 times more likely to die from the infection than girls infected by other girls ^2^ (Figure 6.a). Similarly, boys infected by girls were 1.7 times more likely to die than boys infected by other boys (Figure 6.b). Overall, cross-sex infections with measles were twice as virulent, on average, when compared to average same-sex infections (Figure 6.c). This same qualitative result has been replicated by other studies ^28^ and extended to infections with chickenpox and polio ^25,31^. While these complex patterns of virulence are well established they remain a mystery. Furthermore, differences in virulence between cross-sex and same-sex infections cannot be explained by differences in the immune system of girls and boys alone. The latter would result in different virulence in girls and boys but the lack of effects due to the origin of the infection would result in the cancellation of these differences; if there are only sex effects 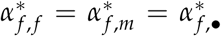 and 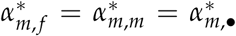, cross-sex and same-sex virulence cannot differ on average.

**Figure 6:**
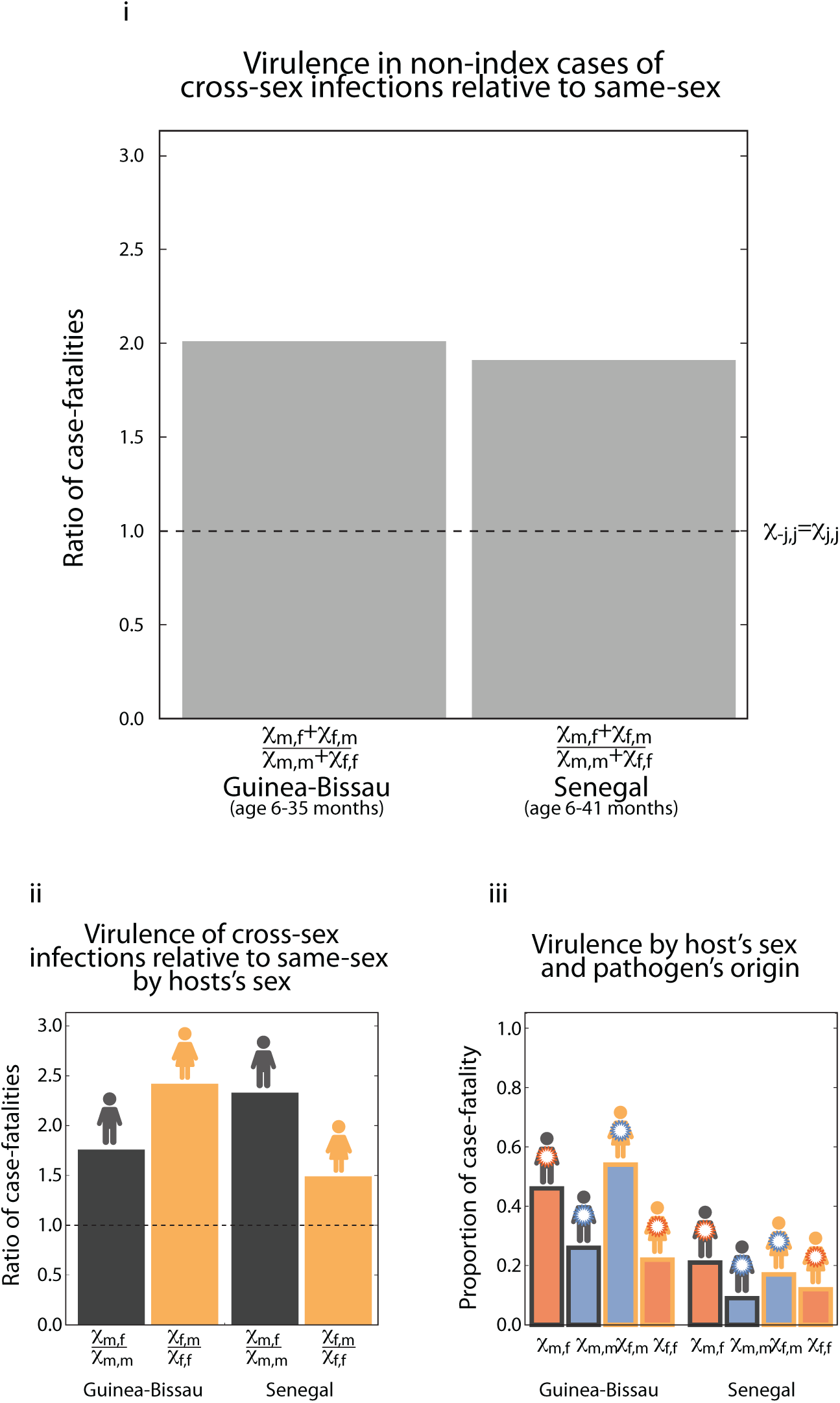
Virulence of cross-sex and same-sex measles infections Panel i: Virulence of cross-sex measles infections relative to same-sex ones. Shows the ratio of case fatalities (*χ*) in cross-sex infections relative to same-sex infections, defined as (*χ*_*m f*_ + *χ*_*f m*_)/(*χ*_*mm*_ + *χ*_*f f*_), in two studies conducted in children 6-35 months old in Guinea-Bissau and Senegal. Notice that in both cases infections from the opposite sex are roughly twice as virulent as infections from the same sex. **Panel ii: Virulence of cross-sex measles infections relative to same-sex ones within the sexes**. Shows the ratio of case fatalities in cross-sex infections relative to same-sex infections broken down by sex of the infected. This ratio in infected boys is given by the proportion of boys infected by girls relative to the one infected by boys, that is *χ*_*m f*_ /*χ*_*mm*_ and in infected girls is given by the proportion of girls infected by boys relative to the one infected by girls, that is *χ*_*f m*_/*χ*_*f f*_. Notice that in all cases infections from the opposite-sex are more virulent than infections from the same-sex, that is *χ¬*_*jj*_/*χ*_*jj*_ *>* 1, ranging between 1.5 and 2.5 times more virulent. **Panel iii: Virulence of measles infections within sexes and origin of the infection**. Shows the proportion of case-fatalities in each of the sexes when infected by each of the other sexes in the two studies conducted in Guinea-Bissau and Senegal ^2,28^.

We argue that epigenetically inherited and acquired marks allow the evolution of sex-&-origin-specific virulence in pathogens in ways that can explain greater virulence of cross-sex infections as opposed to same-sex infections, as observed in measles, chickenpox and polio ^2,25,28,31^. We devised two plausible scenarios in which such complex patterns may evolve; both cases require subpopulations differ with respect to their sex-specific contact patterns. For example, a population where the pathogen is transmitted within two groups of individuals. One group may be formed largely by individuals with primarily same-sex interactions, and the other group by individuals with largely cross-sex interactions. One way in which greater virulence of cross sex-infections may evolve is when, in addition to the contact structure, the two groups differ in their sex-ratio and or size (Figures 3 and 5.b). Alternatively, greater virulence in cross-sex infections may evolve when in addition to the contact structure there is greater same-sex transmissibility (Figure 4 and 5.b).

Our research opens up the intriguing possibility of developing epigenetic therapies by interfering with pathogen’s epigenetic memory. While epigenetic therapies have been widely used in the treatment of cancer ^13,20^, their use in the treatment of infectious diseases is only beginning and focuses on the epigenetic modification of the host genome. Here we propose targeting the pathogen’s epigenome. When pathogens of male origin (resp. female origin) are selected for lower virulence, our model predicts that pathogen’s genes, whose greater expression results in higher virulence (*virulence enhancer*), will be under-expressed when in pathogens of male-origin (resp female-origin) (Figure 7). On the contrary, when pathogens with female-origin (resp female-origin) are selected for higher virulence, our results predict that pathogen’s genes whose greater expression results in lower virulence (*virulence inhibitor*) will be under-expressed when in pathogens of female-origin (resp male-origin) (Figure 7).

**Figure 7:**
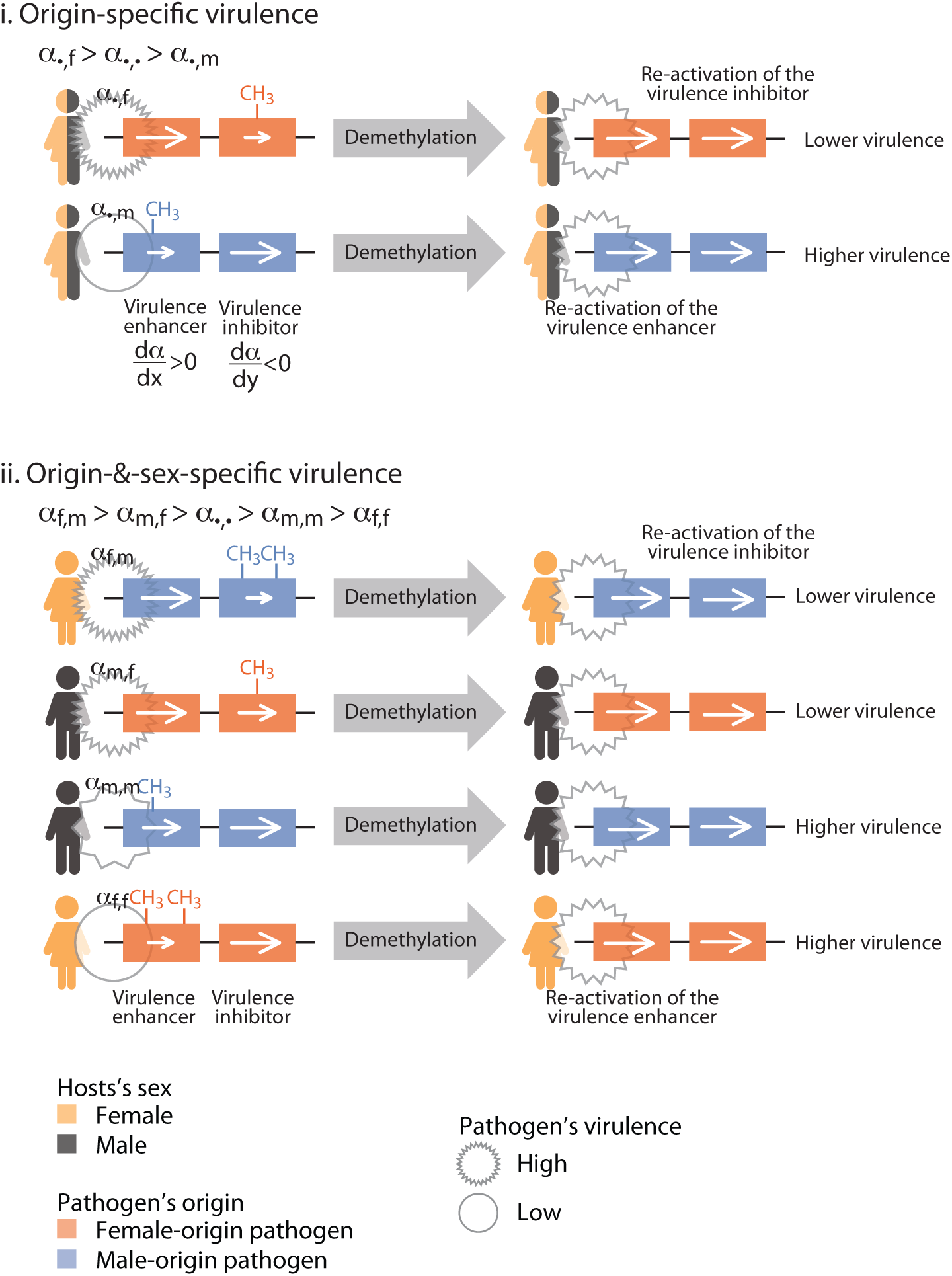
Predictions on the effects of epigenetic therapy on the virulence of an infection. We consider haploid pathogens with two types of genes: one whose greater expression *x* enhances virulence, 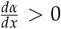, (henceforth *virulence enhancer*), and another one whose greater expression *y* inhibits virulence, 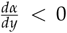, (henceforth *virulence inhibitor*). Epigenetically inherited marks are established as methylation of promoter regions resulting in reduced expression of the gene. Epigenetic interventions consist on de-methylating the pathogen’s genome. **Panel i: On pathogens showing origin-specific virulence**. When female-origin virulence is higher (*α*_•*f*_ *> α*_•*m*_), pathogens of female-origin are selected to methylate the promoter of virulence inhibitor genes thus incrementing the virulence of the infection. We predict that epigenetic interventions consisting in de-methylating infections from females will re-activate virulence inhibitors and thus reduce the virulence of the infection. **Panel ii: On pathogens showing origin-&-sex-specific virulence**. When virulence of pathogens with male-origin in females is higher (*α* _*f m*_ *> α*_*m f*_ *> α*_*mm*_ *> α* _*f f*_), pathogens with male-origin in females are selected to methylate the promoter of virulence inhibitor genes thus incrementing the virulence of the infection. We predict that epigenetic interventions consisting in de-methylating infections in females from females, will re-activate virulence inhibitors and thus reduce the virulence of the infection. Notice however, that pathogens with the same male-origin but infecting females, will respond to the same epigenetic intervention in the opposite manner, that is incrementing the virulence of the infection.

Trans-generational epigenetic marks are often implemented via differential methylation of DNA or histones near the promoter of a gene, although his is not the only mechanism ^12,29^. The epigenetic intervention we propose here is the de-methylation of the pathogen’s genome. This can be achieved through different pharmacological agents, i.e DNA methyltransferase inhibitors ^12,29^. When natural selection favours higher virulence of infections coming from females, we predict that genome-wide demethylation of pathogens originating from females will reduce the virulence of the infection (Figure 7). Similarly, when natural selection favours higher virulence of infections coming from the opposite sex, we predict that genome-wide de-methylation of pathogens originating from males infecting females and originating from females infecting males will reduce the virulence of the infection (Figure 7). We thus propose that it could be possible to reduce the virulence of infections (including measles) by using pharmacological agents that de-methylate the pathogen’s genome (counter-intuitively these will be targeting pathogens genes that down regulate virulence).

More generally, our work suggests that complex patterns of virulence (origin-specific or origin-&-sex specific) can be understood in terms of the selective pressures that affect the pathogen. Attention can not only be paid to biological differences that exist between host types. We propose the intriguing possibility that pathogen’s memories of past environments may be driving the virulence of infections. Our research focuses on the sex of the previous host but our results could be extended to other characters (e.g. the socioeconomic status of hosts).The possibility that there may be a variety of epigenetically inherited memories advocates the need for personalised approaches to infections in which factors like sex or social status can inform the best course of action to treat a disease.

## Methods

See separate file

## Acknowledgements

FU thanks Mateo Úbeda Tran for invaluable advice and support.

